## Supplemental Information for "Do molecular fingerprints identify diverse active drugs in large-scale virtual screening? (no)"

Since, the ROC AUC is not well-suited for evaluating the early recognition of active molecules, metrics such as the Boltzmann-Enhanced Discrimination of the Receiver Operating Characteristic[1] (BEDROC) and the sum of the log of ranks statistic[2, 3] (SLR) have been proposed. To capture early enrichment, we have calculated the Boltzmann-Enhanced Discrimination of the Receiver Operating Characteristic (BEDROC) metric which assigns more weight to early ranked molecules:

$$BEDROC = \frac{\sum_{i=1}^{n} e^{-\alpha r_i/N}}{R_a \left( \frac{1-e^{-\alpha}}{e^{\alpha/N}-1} \right)} \times \frac{R_a \sinh(\alpha/2)}{\cosh(\alpha/2) - \cosh(\alpha/2 - \alpha R_a)} + \frac{1}{1 - e^{\alpha(1-R_a)}} \quad (1)$$

where $n$ is the number of actives, $N$ the total number of compounds, $R_a = n/N$, the ratio of actives to inactives in the dataset and $r_i$, the rank of the $i^{th}$ active. The parameter $\alpha$ is used to emphasize early recognition. The value of $\alpha$ value is chosen so that 0.5% ($\alpha = 321.9$), 2% ($\alpha = 80.5$) or 8% ($\alpha = 20.0$) of the top-ranked molecules account for 80% of the BEDROC score.

We also report the normalized form of the sum of log rank statistic[3] (NSLR):

$$NSLR = \frac{-\sum_{i=1}^{n} \log \frac{r_i}{N}}{-\sum_{i=1}^{n} \log \frac{i}{N}} \quad (2)$$

where $n$ is the number of actives among the $N$ available compounds and $r_i$ is the rank of the $i^{th}$ active. The negative logarithm emphasizes early recognition. The denominator in the equation provides a theoretical maximum when a VS method ranks all actives within the first $n$ positions. NSLR varies between 0 and 1, where the latter is the best achievable ranking.

| FP | DEKOIS | DUDE | MUV | LIT-PCBA | DEKOIS | DUDE | MUV | LIT-PCBA |
|---|---|---|---|---|---|---|---|---|
| | | **AUC_S** | | | | **AUC_L** | | |
| AVALON | 0.72 | 0.73 | 0.60 | 0.55 | 0.70 | 0.77 | 0.60 | 0.55 |
| ECFP0 | 0.70 | 0.77 | 0.53 | 0.50 | 0.70 | 0.77 | 0.53 | 0.50 |
| ECFP2 | 0.77 | 0.81 | 0.54 | 0.51 | 0.77 | 0.81 | 0.54 | 0.51 |
| ECFP4 | 0.76 | 0.80 | 0.54 | 0.51 | 0.77 | 0.81 | 0.54 | 0.51 |
| ECFP6 | 0.75 | 0.78 | 0.54 | 0.52 | 0.77 | 0.80 | 0.54 | 0.51 |
| FCFP0 | 0.66 | 0.69 | 0.54 | 0.52 | 0.66 | 0.69 | 0.54 | 0.52 |
| FCFP2 | 0.76 | 0.75 | 0.55 | 0.52 | 0.76 | 0.75 | 0.55 | 0.51 |
| FCFP4 | 0.78 | 0.76 | 0.54 | 0.52 | 0.78 | 0.76 | 0.55 | 0.51 |
| FCFP6 | 0.78 | 0.75 | 0.54 | 0.52 | 0.78 | 0.76 | 0.54 | 0.51 |
| | | **DRF_S** | | | | **DRF_L** | | |
| AVALON | 0.30 | 0.18 | 0.85 | 0.97 | 0.25 | 0.16 | 0.89 | 0.98 |
| ECFP0 | 0.33 | 0.13 | 0.97 | 0.96 | 0.33 | 0.13 | 0.97 | 0.96 |
| ECFP2 | 0.19 | 0.09 | 0.99 | 1.01 | 0.18 | 0.08 | 0.96 | 0.98 |
| ECFP4 | 0.19 | 0.09 | 0.99 | 1.00 | 0.18 | 0.09 | 0.99 | 0.99 |
| ECFP6 | 0.20 | 0.10 | 0.99 | 0.98 | 0.19 | 0.09 | 0.99 | 1.01 |
| FCFP0 | 0.35 | 0.23 | 0.41 | 0.42 | 0.35 | 0.23 | 0.41 | 0.42 |
| FCFP2 | 0.24 | 0.19 | 0.93 | 1.01 | 0.24 | 0.19 | 0.93 | 1.00 |
| FCFP4 | 0.20 | 0.15 | 0.93 | 1.01 | 0.20 | 0.16 | 0.93 | 1.01 |
| FCFP6 | 0.20 | 0.15 | 0.96 | 0.98 | 0.20 | 0.16 | 0.94 | 1.02 |

*Table S1: Summary of the VS screening performances using extended length fingerprints. The impact of fingerprint length on the VS metrics was assessed using the default ("_S") length (1024 bits) and the long ("_L") form (16384 bits).*

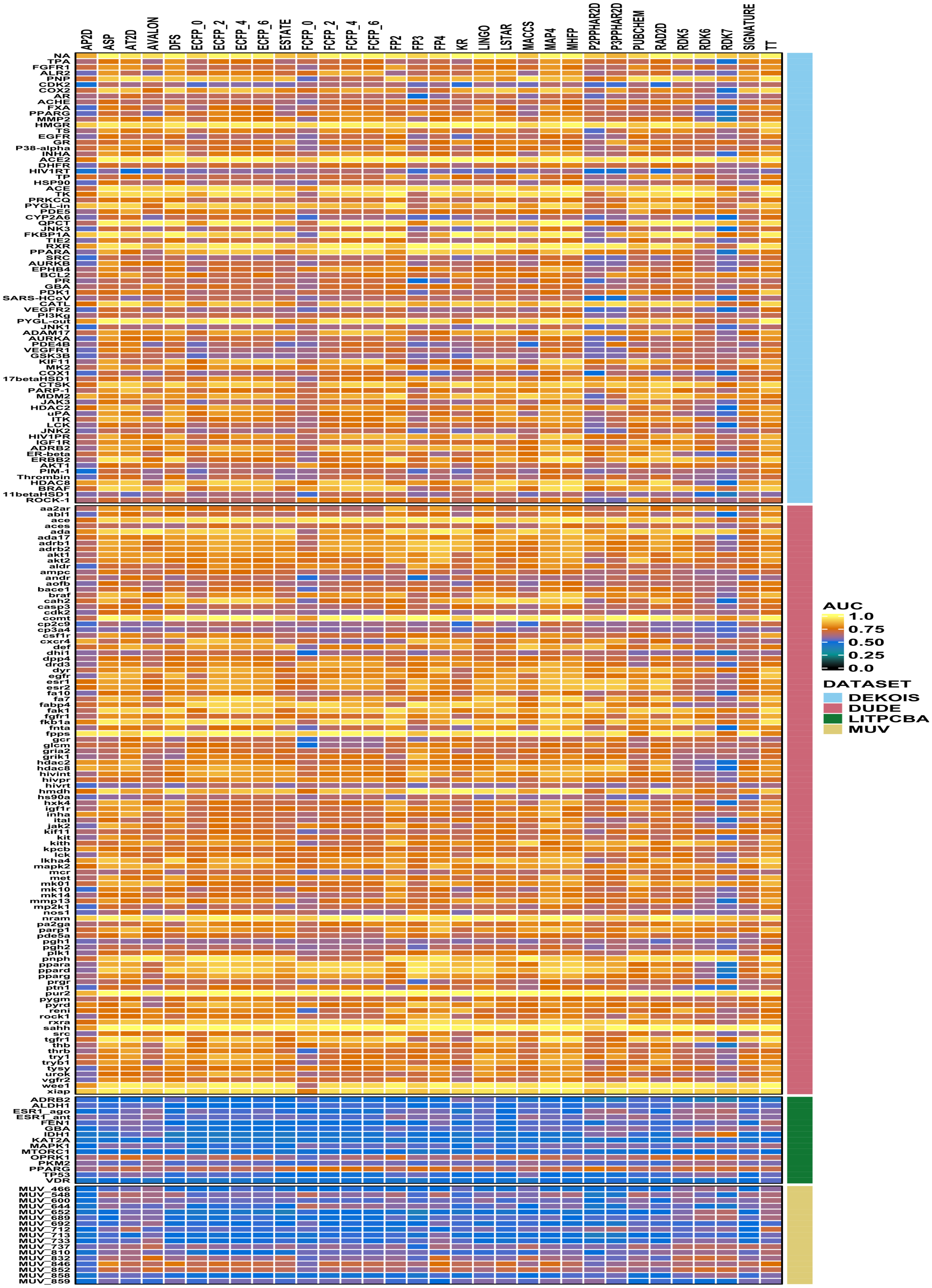


Figure F1: Heatmap of the area under the curve (AUC) obtained by the different fingerprints for the targets in the benchmark datasets.

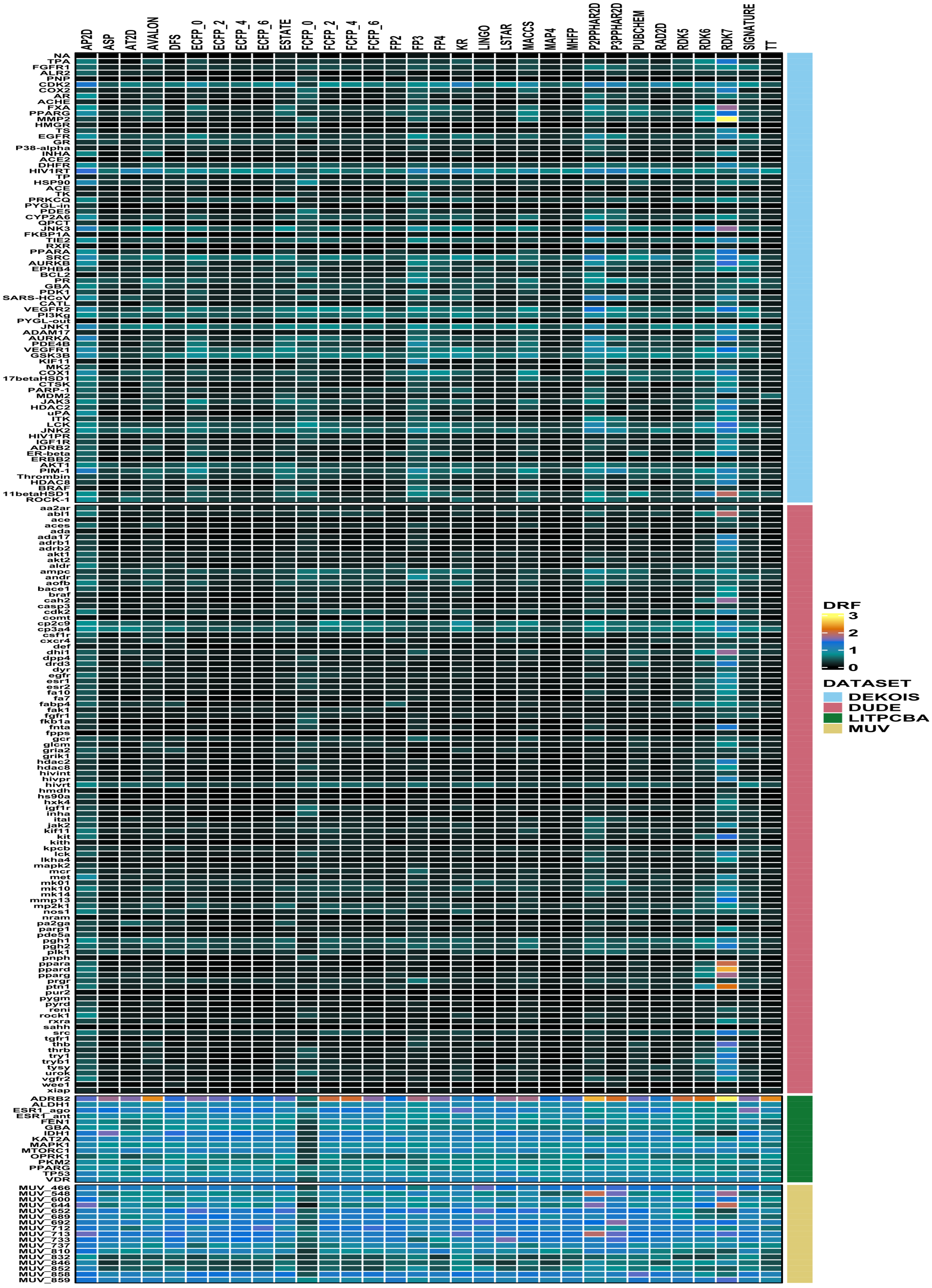


Figure F2: Heatmap of the decoy retention factor ($DRF_{0.1}$) obtained by the different fingerprints for the targets in the benchmark datasets.

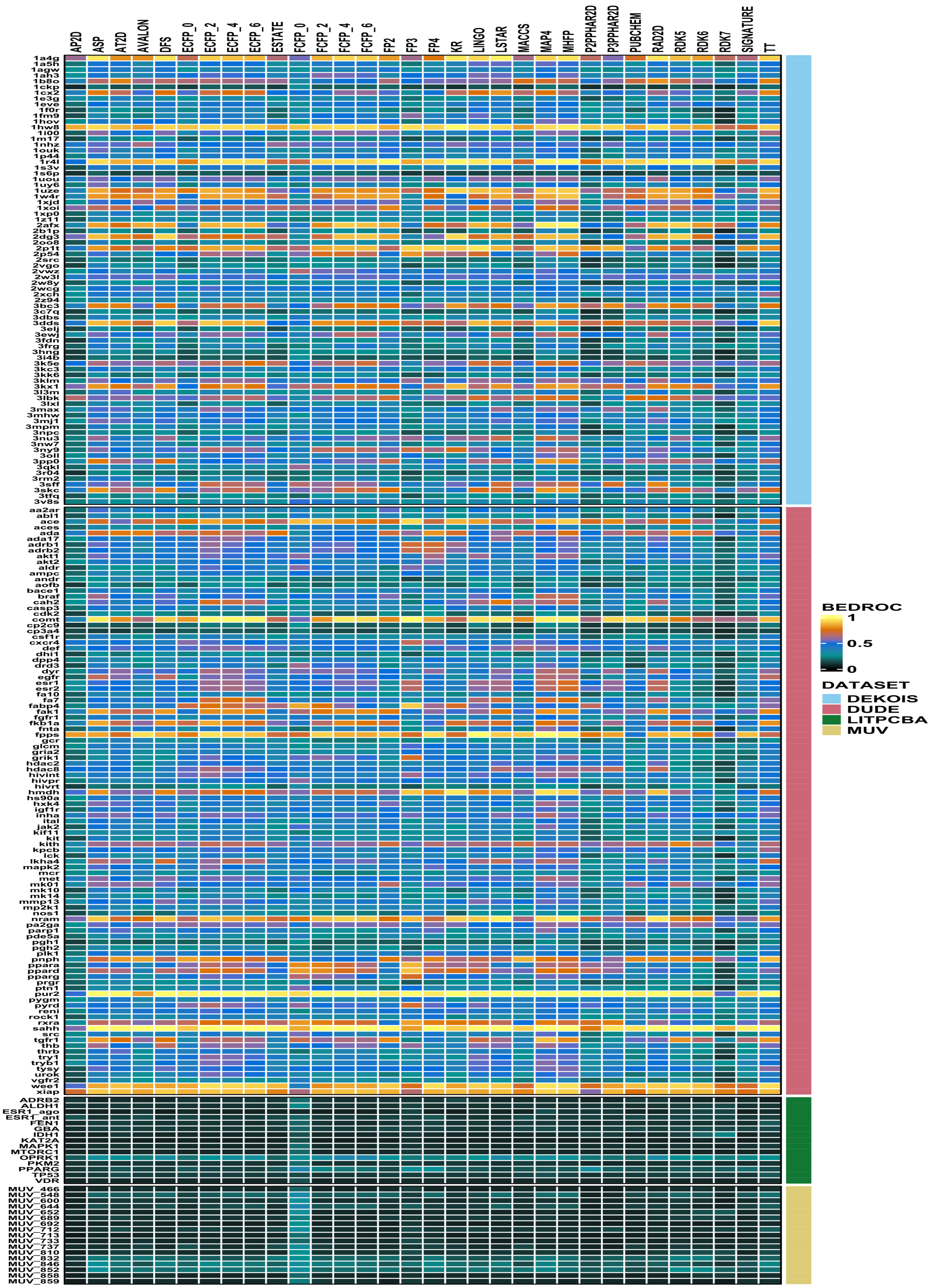


Figure F3: Heatmap of the BEDROC ($\alpha = 20$) values obtained by the different fingerprints for the targets in the benchmark datasets.

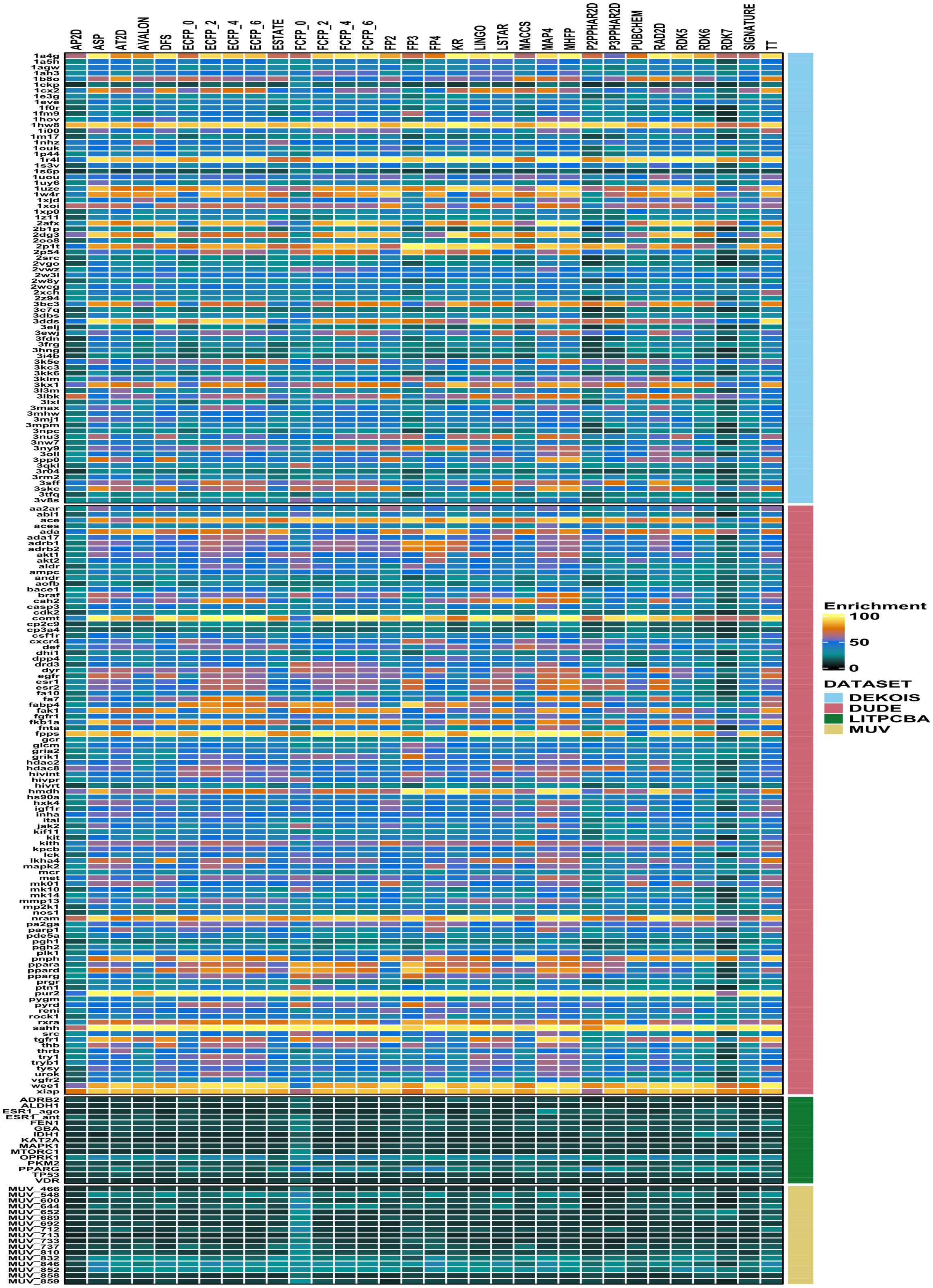


Figure F4: Heatmap of the enrichment factor (top 8%) obtained by the different fingerprints for the targets in the benchmark datasets.

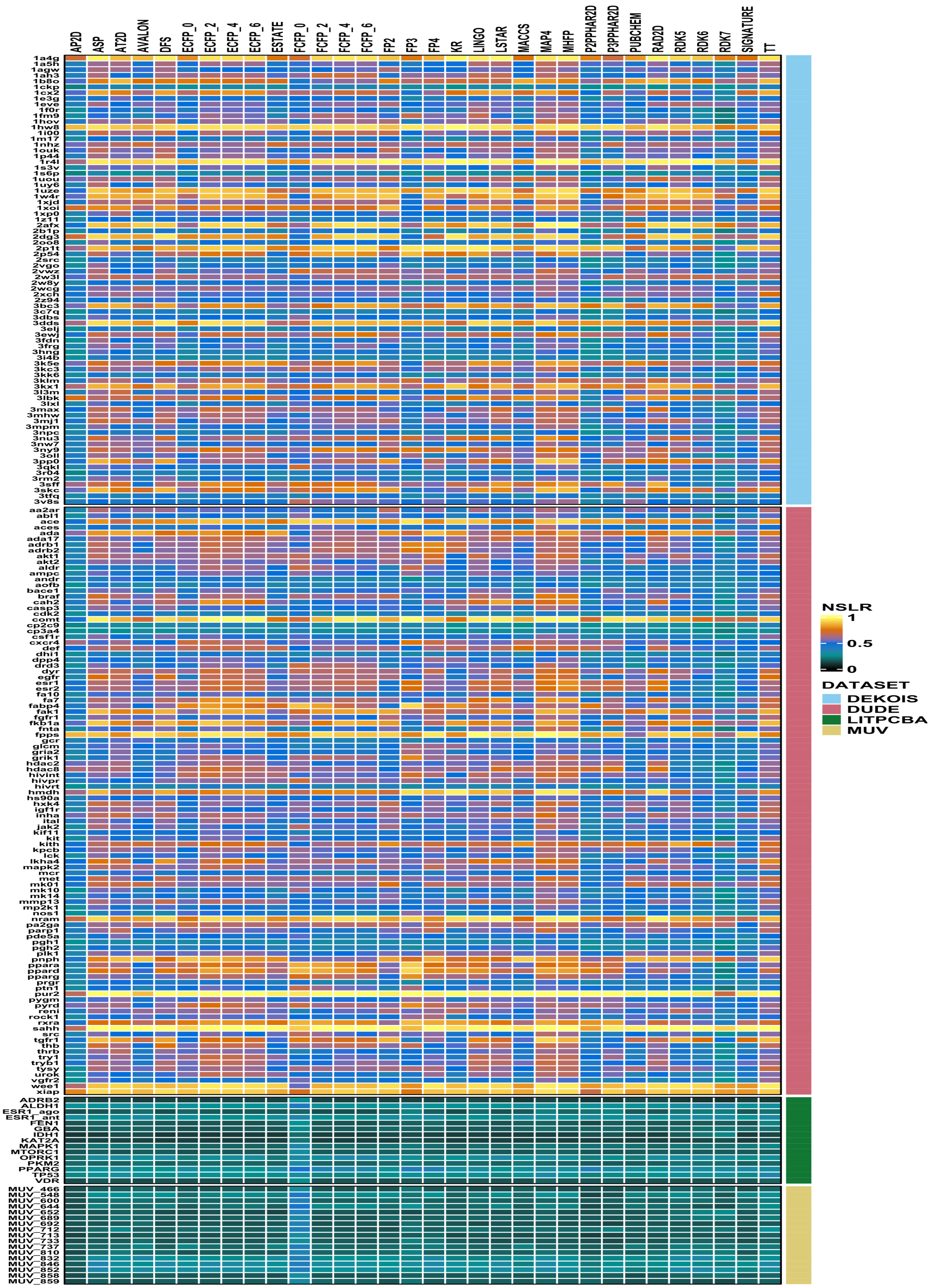

Figure F5: Heatmap of the NSLR values obtained by the different fingerprints for the targets in the benchmark datasets.

PPARG

Active Query
CN(CCOc1ccc(CC(C([N-]2)=O)SC2=O)cc1)c1ncccc1

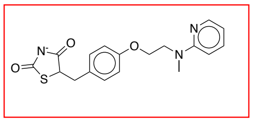

Active hit with Tanimoto >=0.5 (ECFP4)

CCc1cnc(CCOc2ccc(CC(C([N-]3)=O)SC3=O)cc2)cc1

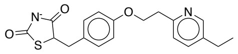

Same scaffold; non-obvious

Figure F6: In the LIT-PCBA analysis, early enrichment was observed for GBA – for all Tanimoto coefficients $t > 0.2$, the fraction of actives with Tanimoto score $> t$ is much larger than the fraction of decoys with that score (see Figure 2 in the main text). There are 24 actives in the OPRK1 data set. We manually inspected the structures of the 1 compound with Tanimoto score $> 0.5$ to the initial query. This compound is built on the same scaffold as the query (in red), but is a non-obvious variant on that scaffold.

GBA

Active Query

Cc(cc1)cc(Cl)c1NC(CN(c1cc(C)cc(C)c1)S(c(c(C)nc([O-])n1)c1[O-])(=O)=O)=O

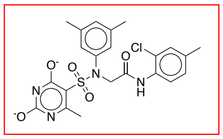

Active hits with Tanimoto >=0.5 (ECFP4)

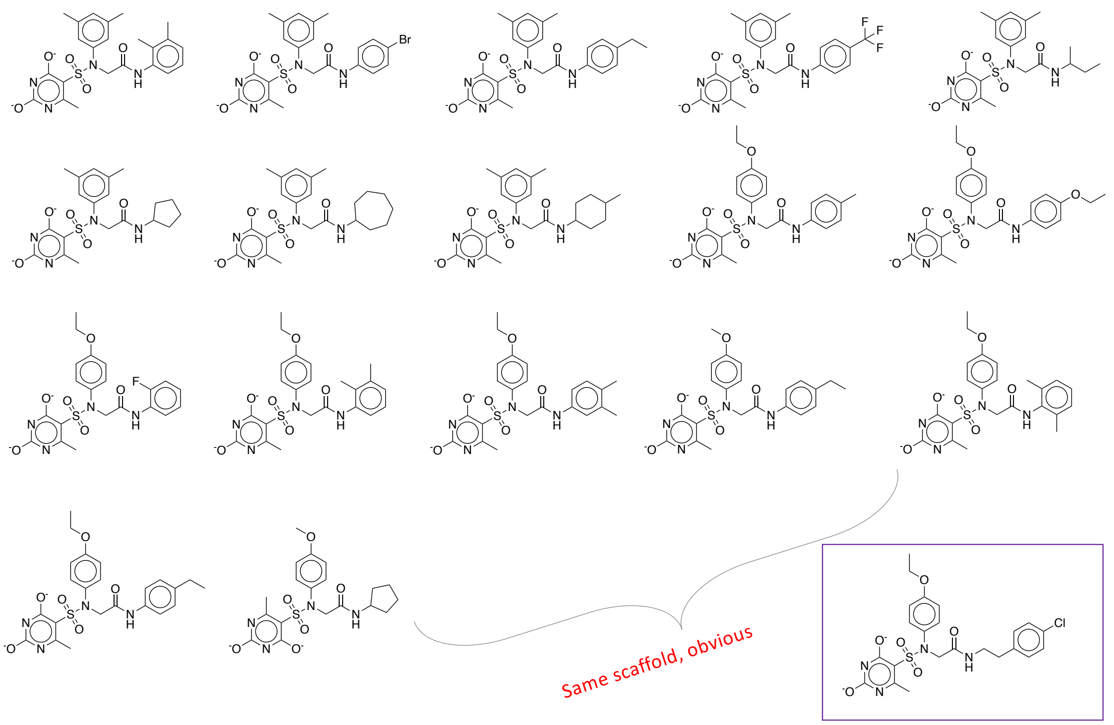


Same scaffold; non-obvious.

Cc1cc(N(CC(Nc2c(C)c(C)ccc2)=O)S(c(c(C)nc([O-])n2)c2[O-])(=O)=O)cc(C)c1
Cc1cc(N(CC(Nc(cc2)ccc2Br)=O)S(c(c(C)nc([O-])n2)c2[O-])(=O)=O)cc(C)c1
CCc(cc1)ccc1NC(CN(c1cc(C)cc(C)c1)S(c(c(C)nc([O-])n1)c1[O-])(=O)=O)=O
Cc1cc(N(CC(Nc2ccc(C(F)(F)F)cc2)=O)S(c(c(C)nc([O-])n2)c2[O-])(=O)=O)cc(C)c1
CCC(C)NC(CN(c1cc(C)cc(C)c1)S(c(c(C)nc([O-])n1)c1[O-])(=O)=O)=O
Cc1cc(N(CC(NC2CCCC2)=O)S(c(c(C)nc([O-])n2)c2[O-])(=O)=O)cc(C)c1
Cc1cc(N(CC(NC2CCCCCC2)=O)S(c(c(C)nc([O-])n2)c2[O-])(=O)=O)cc(C)c1
CC(CC1)CCC1NC(CN(c1cc(C)cc(C)c1)S(c(c(C)nc([O-])n1)c1[O-])(=O)=O)=O
CCOc(cc1)ccc1N(CC(Nc1ccc(C)cc1)=O)S(c(c(C)nc([O-])n1)c1[O-])(=O)=O
CCOc(cc1)ccc1NC(CN(c(cc1)ccc1OCC)S(c(c(C)nc([O-])n1)c1[O-])(=O)=O)=O
CCOc(cc1)ccc1N(CC(Nc(cccc1)c1F)=O)S(c(c(C)nc([O-])n1)c1[O-])(=O)=O
CCOc(cc1)ccc1N(CC(Nc1c(C)c(C)ccc1)=O)S(c(c(C)nc([O-])n1)c1[O-])(=O)=O
CCOc(cc1)ccc1N(CC(Nc1cc(C)c(C)cc1)=O)S(c(c(C)nc([O-])n1)c1[O-])(=O)=O
CCc(cc1)ccc1NC(CN(c(cc1)ccc1OC)S(c(c(C)nc([O-])n1)c1[O-])(=O)=O)=O
CCOc(cc1)ccc1N(CC(Nc1c(C)cccc1C)=O)S(c(c(C)nc([O-])n1)c1[O-])(=O)=O
CCc(cc1)ccc1NC(CN(c(cc1)ccc1OCC)S(c(c(C)nc([O-])n1)c1[O-])(=O)=O)=O
Cc1nc([O-])nc([O-])c1S(N(CC(NC1CCCC1)=O)c(cc1)ccc1OC)(=O)=O
CCOc(cc1)ccc1N(CC(NCCc(cc1)ccc1Cl)=O)S(c(c(C)nc([O-])n1)c1[O-])(=O)=O

Figure F7: In the LIT-PCBA analysis, early enrichment was observed for GBA – for all Tanimoto coefficients $t > 0.2$, the fraction of actives with Tanimoto score $> t$ is much larger than the fraction of decoys with that score (see Figure ??). There are 163 actives in the OPRK1 data set. We manually inspected the structures of the 18 compounds with Tanimoto score $> 0.5$ to the initial query. All 18 appear are built on the same scaffold as the query (in red), and all but one is an obvious variation that should be identified through standard enumeration (i.e. no new scaffold are explored).
